# Computational docking reveals evolutionary conservation of a specific interaction between 15d-Prostaglandin-J2 and eIF4A

**DOI:** 10.1101/237610

**Authors:** So Jeong Yun, Hyunjoon Kim, Seung Gee Lee, Seung-Hyun Jung, Joon Hyun Kim, Jeong Eun Ryu, N. Jiten Singh, Jouhyun Jeon, Jin-Kwan Han, Cheol-Hee Kim, Sanguk Kim, Kwang S. Kim, Sung Key Jang, Woo Jae Kim

## Abstract

15-deoxy-delta 12,14-prostaglandin J2 (15d-PGJ2) is anti-inflammatory/antineoplastic prostaglandin which functions through covalent binding to cysteine residues of various target proteins. We previously showed that 15d-PGJ2 mediated anti-inflammatory responses are dependent on the translational inhibition through its interaction with eIF4A. Binding of 15d-PGJ2 to eIF4A specifically blocks the interaction between eIF4G and eIF4A leads to the formation of stress granules (SGs), which cluster mRNAs with inhibited translation. Here we show that the binding between 15d-PGJ2 and eIF4A specifically blocks the interaction between the MIF4G domain of eIF4G and eIF4A. To reveal the mechanism of this interaction, we used computational simulation-based docking studies and identified that the carboxyl tail of 15d-PGJ2 could stabilize the binding of 15d-PGJ2 to eIF4A through arginine 295 of eIF4A, which is the first suggestion that the 15d-PGJ2 tail play a physiological role. Interestingly, the putative 15d-PGJ2 binding site on eiF4A is conserved across many species, suggesting a biological role. Our data propose that studying 15d-PGJ2 and its targets will may uncover new therapeutic approaches in anti-inflammatory drug discovery.

## INTRODUCTION

15-deoxy-delta 12,14-prostaglandin J2 (15d-PGJ2) is an anti-inflammatory and antineoplastic prostaglandin. Although 15d-PGJ2 is known as an agonist of peroxisome proliferator-activated receptor-gamma (PPARγ), which is a transcriptional modulator that represses transcription of pro-inflammatory mRNAs, evidence suggests that 15d-PGJ2 also can function independently of PPARγ (Nosjean and Boutin, 2002). It has been reported that the PPARγ-independent action of 15d-PGJ2 resulted from the covalent modification of cysteine residues of target proteins. For example, 15d-PGJ2 blocks pro-inflammatory NF-κB signaling cascades independently of PPARγ through direct interactions with signaling molecules such IKK (IκB kinase) (Straus et al., 2000). Other physiological activities of 15d-PGJ2, such as cytoprotection and inhibition of cell proliferation, have also been reported that occurred through this direct binding property of 15d-PGJ2 (Pereira et al., 2006). 15d-PGJ2 is a member of the cyclopentone-type prostaglandins (PGs). Cyclopentone-type PGs, unlike other classes of PGs, contain an electrophilic α,ß-unsaturated ketone moiety in the cyclopentenone ring. This reactive center of the cyclopentone PGs can act as a Michael addition acceptor and react with nucleophiles, such as the free thiol group of the glutathione and cysteine residues in cellular proteins. These properties of 15d-PGJ2 could explain the biological activities of 15d-PGJ2 independent of PPARγ (Kondo et al., 2002; Shibata, 2015).

Among the cellular proteins that are covalently modified by 15d-PGJ2, eIF4A is the only factor that directly modulates the initiation of translation (Kim et al., 2007). eIF4A is the founding member of the “DEAD-box” family of ATP-dependent helicases (Oberer et al., 2005; Rogers et al., 2001). It consists of two distinct domains connected through a short linker, and both domains are required for proper function of the helicase. The ATPase activity of eIF4A is stimulated by eIF4G, and the helicase activity of eIF4A either alone or as part of the eIF4F complex is stimulated by eIF4B (Rozen et al., 1990; Schütz et al., 2008)

Recent studies focus on the function of eIF4A and its relation to cancer and inflammation for the following reasons. First, it was reported that PDCD4, a novel tumour suppressor protein, interacts with eIF4A, which results in the inhibition of helicase activity and translation (Yang et al., 2003), indicating that blocking the cell-proliferative function of eIF4A is a critical step to suppress tumorigenesis. Second, pateamine A (PatA), a potent anti-proliferative and pro-apoptotic marine natural product, can bind to and enhance the intrinsic enzymatic activities of eIF4A. PatA inhibits the eIF4A-eIF4G association and promotes the formation of a stable ternary complex between eIF4A and eIF4B (Low et al., 2005). Finally, our previous report suggests that 15d-PGJ2 covalently binds to a cysteine residue (C264) in eIF4A, resulting in the inhibition of translation initiation and formation of stress granules (SGs) (Kim et al., 2007). Following our previous results, here we report further characterization of the interaction between 15d-PGJ2 and eIF4A. Also, we will show the effect of 15d-PGJ2 on various model organisms following our findings on evolutionary conserved 15d-PGJ2 binding sites of eIF4A across species.

## RESULTS AND DISCUSSION

### 15d-PGJ2 binding to eIF4A specifically blocks the interaction between MIF4G domain of eIF4G and eIF4A

We previously reported that the direct interaction between 15d-PGJ2 and eIF4A can specifically block the eIF4A-eIF4G binding and inhibit translation initiation (Kim et al., 2007). To further analyze this interaction, we performed a series of immunoprecipitation experiments using other eIF4GI interacting proteins. When we immunoprecipitated FLAG tagged eIF4A1, eIF4E or eIF3c then performed immunoblotting with eIF4G antibody, we identified that the interactions of eIF4G with eIF4E (Fig. 1A, lanes 3 and 4) was partially affected by 15d-PGJ2 while association of eIF4G with eIF3c was not affected (Fig. 1A, lanes 5 and 6). The interaction between eIF4A1 and eIF4GI is blocked by 15d-PGJ2 as we previously described (Fig. 1A, lanes 1 and 2) (Kim et al., 2007). We also confirmed that the RNA-mediated interaction between eIF4A and PABP is not inhibited by 15d-PGJ2 treatment, rather 15d-PGJ2 enhance the RNA-mediated interaction between eIF4A and PABP (Fig. S1A). This data is consistent with our previous report that the RNA-binding activity of eIF4A is increased when it binds to 15d-PGJ2 (Kim et al., 2007). In addition, the interaction between eIF4A and eIF4B is not affected by 15d-PGJ2 binding to eIF4A (Fig. S1B). These data suggest that 15d-PGJ2 binding to eIF4A specifically blocks the interaction between eIF4G and eIF4A while promoting its binding to PABP.

**Figure 1.**
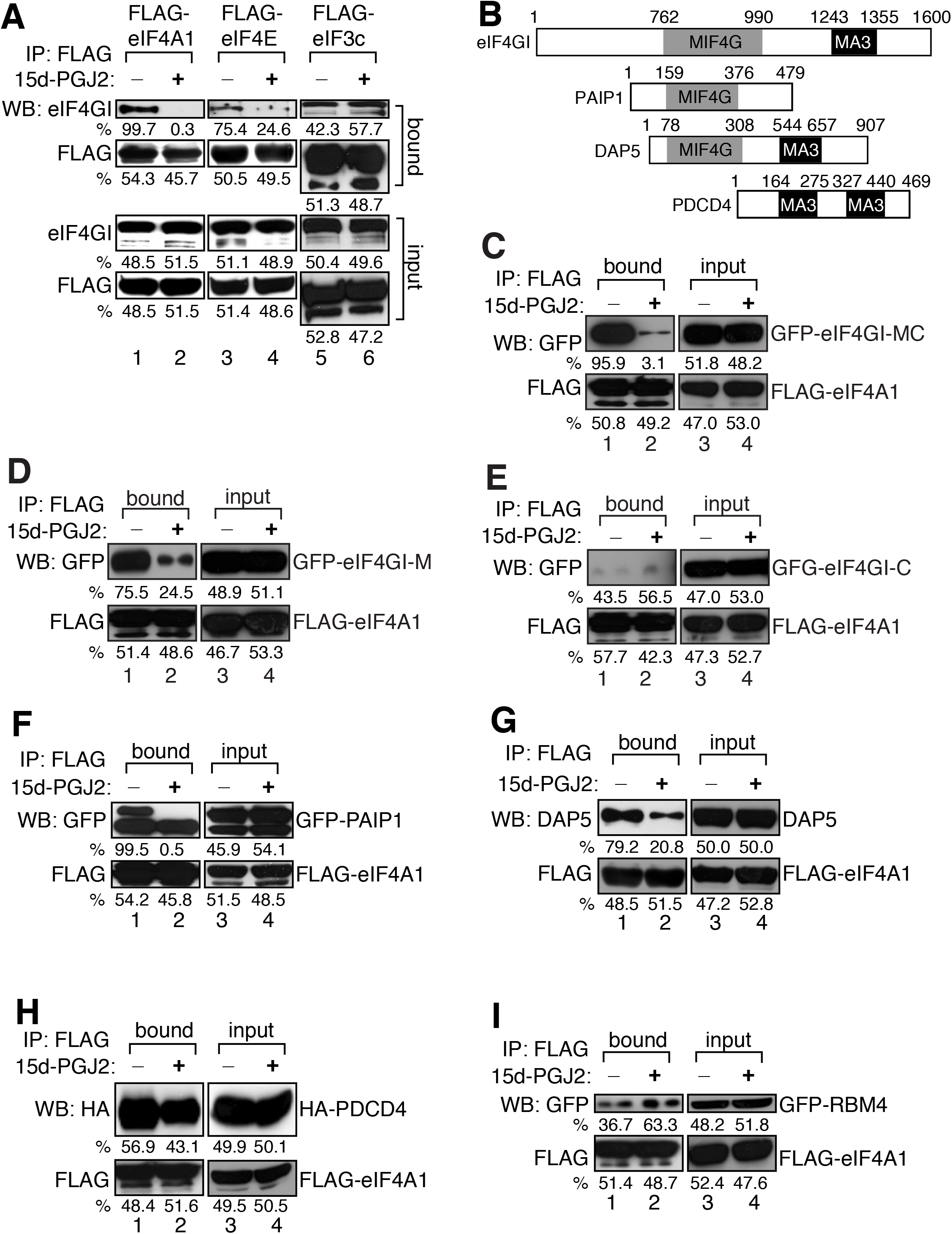
15d-PGJ2 blocks the interaction between MIF4G domain of eIF4G with eIF4A. (A) 293T cells were transfected with FLAG-eIF4A1 (lanes 1 and 2), FLAG-eIF4E (lanes 3 and 4), or FLAG-eIF3c (lanes 5 and 6). Cells were lysed then treated with EtOH or 10 μM of 15d-PGJ2 at 30°C for 1 hour. Immunoprecipitation was performed using anti-FLAG antibody then Western-blot analysis was performed with anti-FLAG and eIF4GI antibodies. The quantification of the relative band intensity was performed using ImageJ (See **EXPERIMENTAL PROCEDURES**). Thus, the numbers described below the band indicates the relative percentage of band intensity compared to neighboring band. If the intensity of neighboring bands is exactly same, the values will be 50:50. The same quantification methods are used in all figures unless specified differently. (B) eIF4A binding domain structures of eIF4GI homologues were illustrated based on the Pfam graphical view of domain structure (http://pfam.sanger.ac.uk). MIF4G domain homologues were marked as grey, MA3 domain homologues as black. (C) 293T cells were co-transfected with GFP-eIF4GI-MC and FLAG-eIF4A1. Cells were lysed then treated with EtOH or 10 μM of 15d-PGJ2 at 30°C for 1 hour. Immunoprecipitation was performed as described above then Western-blot analysis was performed with anti-FLAG and anti-GFP antibodies. (D) 293T cells were co-transfected with GFP-eIF4GI-M and FLAG-eIF4A1. Cells were lysed then treated with EtOH or 10 μM of 15d-PGJ2 at 30°C for 1 hour. Immunoprecipitation was performed as described above then Western-blot analysis was performed with anti-FLAG and anti-GFP antibodies. (E) 293T cells were co-transfected with GFP-eIF4GI-C and FLAG-eIF4A1. Cells were lysed then treated with EtOH or 10 μM of 15d-PGJ2 at 30°C for 1 hour. Immunoprecipitation was performed as described above then Western-blot analysis was performed with anti-FLAG and anti-GFP antibodies. (F) 293T cells were co-transfected with GFP-PAIP1 and FLAG-eIF4A1. Cells were lysed then treated with EtOH or 10 μM of 15d-PGJ2 at 30°C for 1 hour. Immunoprecipitation was performed with an anti-FLAG antibody. Western-blot analysis was performed with anti-FLAG and anti-GFP antibodies. (G) 293T cells were transfected with FLAG-eIF4A1. Cells were lysed then treated with EtOH or 10 μM of 15d-PGJ2 at 30°C for 1 hour. Immunoprecipitation was performed as described above then Western-blot analysis was performed with anti-FLAG and anti-DAP5 antibodies. (H) 293T cells were co-transfected with HA-PDCD4 and FLAG-eIF4A1. Cells were lysed then treated with EtOH or 10 μM of 15d-PGJ2 at 30°C for 1 hour. Immunoprecipitation was performed as described above then Western-blot analysis was performed with anti-FLAG and anti-HA antibodies. (I) 293T cells were co-transfected with GFP-RBM4 and FLAG-eIF4A1. Cells were lysed then treated with EtOH or 10 μM of 15d-PGJ2 at 30°C for 1 hour. Immunoprecipitation was performed as described above then Western-blot analysis was performed with anti-FLAG and anti-GFP antibodies.

It has been known that human eIF4G has two domains, MIF4G (HEAT-1) and MA-3 (HEAT-2) for the interaction with eIF4A (Craig et al., 1998; Imataka and Sonenberg, 1997; Lomakin et al., 2000; Marintchev et al., 2009; Oberer et al., 2005; Schütz et al., 2008) (Fig. 1B). However, it is unknown whether eIF4A interacts with two binding domains of eIF4G through its same surface or different surfaces. To identify which eIF4A interaction domain of eIF4GI is sensitive to 15d-PGJ2’s binding to eIF4A, we expressed subdomains of GFP-tagged eIF4GI with FLAG-eIF4A then performed immunoprecipitation assay using anti-FLAG antibody. The interaction between full-length GFP-eIF4GI and FLAG-eIF4A is inhibited by 15d-PGJ2 treatment as expected (Fig. S1C). The interaction between FLAG-eIF4A and GFP-eIF4GI-MC which contains both MIF4G and MA3 domain, was significantly inhibited by 15d-PGJ2 (Fig. 1C). When the GFP-eIF4GI-M, which contains only MIF4G domain was expressed with FLAG-eIF4A, their interaction was also interrupted by 15d-PGJ2 treatment (Fig. 1D). However, the interaction between FLAG-eIF4A and GFP-eIF4GI-C, which contains only MA3 domain was not affected by 15d-PGJ2 (Fig. 1E). We also tested the effect of 15d-PGJ2 on the interaction between eIF4A and eIF4GII, a paralogue of eIF4GI. However, we could not detect a strong interaction between overexpressed GFP-eIF4GII and FLAG-eIF4A nor the effect of 15d-PGJ2 on this interaction (Fig. S1D). We also confirmed that the interactions between eIF4A and the binding domains of eIF4GII are not affected by 15d-PGJ2 (Fig. S1E-G). These data suggest that the interaction between the MIG4G domain of eIF4GI, not eIF4GII and eIF4A is more sensitive to 15d-PGJ2 binding to eIF4A.

To further characterize whether 15d-PGJ2 blocks binding of eIF4A to interactors other than eIF4G, we tested the effect of 15d-PGJ2 on eIF4G homologues containing eIF4A binding domains or on other eIF4A binding partners containing MIF4G or MA3 domains (Fig. 1B). The interactions of eIF4A with PAIP1 or with DAP5, both of which contain regions similar to the MIF4G domain were significantly reduced by 15d-PGJ2 (Fig. 1F and G). The interaction of PDCD4, an MA-3 domain containing protein, with eIF4A was slightly affected by 15d-PGJ2 (Fig. 1H). It has been reported that eIF4A interacts with the RNA binding protein RMB4, which does not contain MIF4G or MA3 domains (Lin et al., 2007). The interaction between RBM4 and eIF4A is not inhibited by 15d-PGJ2, rather their interaction was slightly increased when 15d-PGJ2 was added to the binding reaction (Fig. 1I). All these data suggest that 15d-PGJ2 binding to eIF4A specifically block the interaction between MIF4G domain and eIF4A.

To further investigate the 15d-PGJ2 interacting residues within eIF4A, we decided to use a computational approach to simulate the interaction between 15d-PGJ2 and the structural model of human eIF4A. Homology modeling, which is template-based modeling, constructs an atomic-resolution model structure of the “target” protein using its amino acid sequence and its homologous protein structure (“template”) obtained from experiments. Homology modeling assumes that the protein structures are more conserved than protein sequences. Practically, the proteins with sequence similarities of more than 30% can be used as templates (Yang and Honig, 2000)(Yun, 2011). The method provides accurate models of protein structures, which can be used for the studies of protein-protein and protein-ligand docking, of site-directed mutagenesis, and of catalytic mechanism investigation. Docking simulation predicts the orientation of the binding of small molecules (ligands and drug candidates) to their target proteins and infers the affinity and activity of the small molecules. Therefore, it has played an important role in the rational design of drugs (structure based drug screening). We take advantage of using docking simulation since it samples the conformations of ligands in the binding site of proteins and provides reliable binding modes through assessing the conformations using a scoring function (Vieth et al., 1998).

We used the model structure of 15d-PGJ2 based on a previous study (Fig. 2A, see Methods for details) (Pande and Ramos, 2005). Then we built the model structure of human eIF4A-1 based on the crystal structure of MjDEAD from the *hyperthermophile Methanococccus jannaschii* (PDB id; 1HV8) (see Methods). The sequence homology between MjDEAD and eIF4A-1 was 33.8% and similarity was 54.4 %. We confirmed that nearly all motifs characterizing the DEAD box helicases in eIF4A were conserved in MjDEAD. When we performed the docking simulation, we found that there are 9 plausible residues of eIF4A that might interact with 15d-PGJ2 (E257, D261, T262, C264, D265, R295, L400, D404, I406), which are presented as Van der Waals contact surfaces (Fig. 2C). It is already known that 15d-PGJ2 contains a reactive α,β-unsaturated ketone in the cyclopentenone ring in which an electrophilic carbon is susceptible for Michael addition (Straus and Glass, 2001). Among those amino acid residues of eIF4A that simulation predicted to interact with 15d-PGJ2, only C264 is in proximity to the electrophilic carbon in the head region of 15d-PGJ2 (distance 3.8 Å), which is a distance compatible with covalent bonding to undergo a Michael addition to eIF4A (Fig. 2C). We also confirmed that C264 is located at the most solvent accessible surface among all Cys residues of eIF4A (Fig. 2B), further suggesting C264 is the likely site of modification with 15d-PGJ2 as we previously reported (Kim et al., 2007).

**Figure 2.**
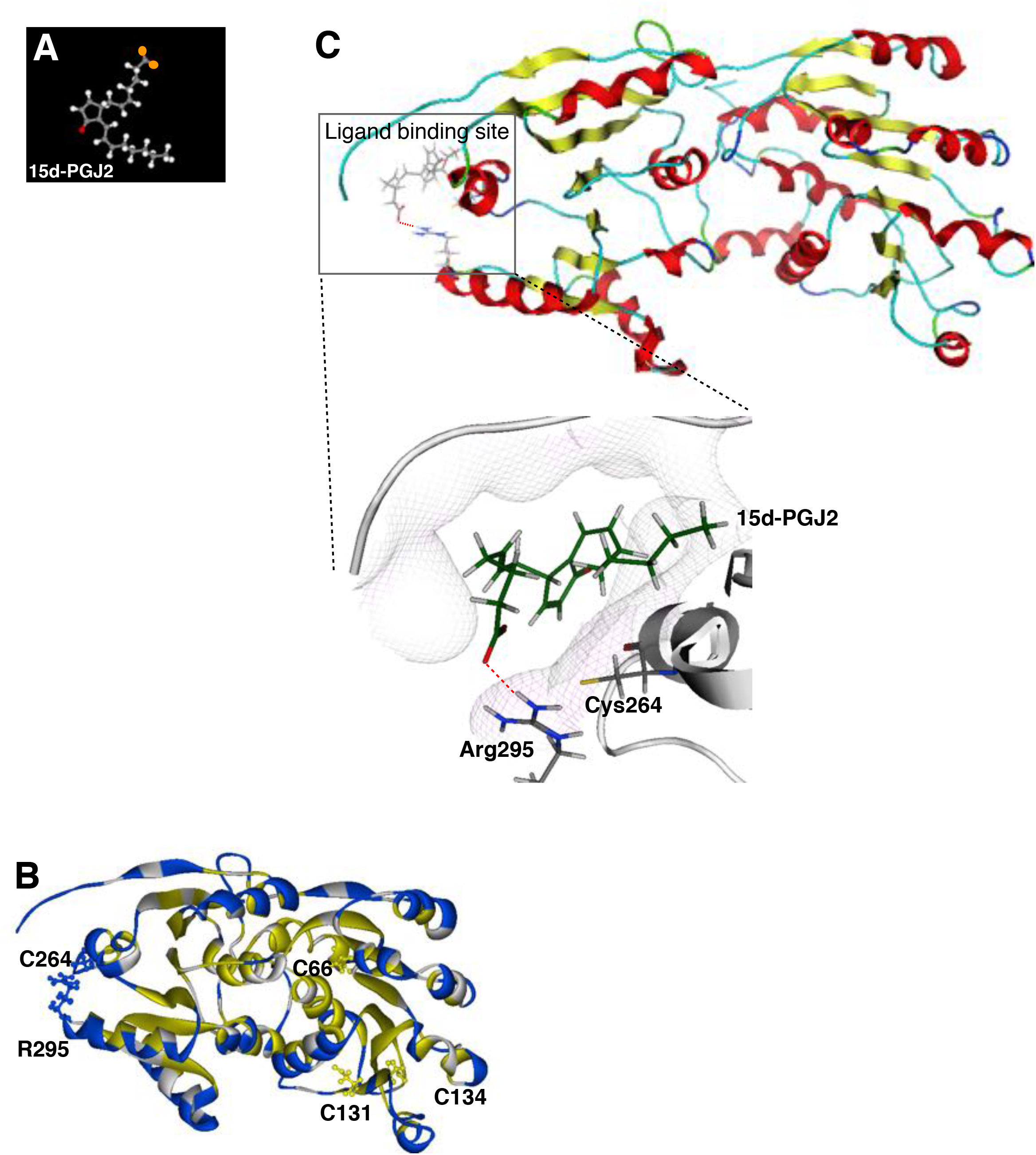
Carboxyl tail of 15d-PGJ2 interacts with R295 of eIF4A in docking simulation. (A) Structure of 15d-PGJ2. The head region of 15d-PGJ2 contains the reactive α,β-unsaturated ketone structure labeled as red color. The carboxyl terminal of tail region also labeled as orange color. (B) Homology model of human eIF4A-1 based on the crystal structure of MjDEAD (PDB id: 1HV8). The Cys residues of eIF4A are marked. Note that only C264 is located in the hydrophilic region (blue), the other Cys residues are in the hydrophobic region (yellow). The yellow colored ribbon indicates hydrophobic (inner) region of protein and blue colored ribbon hydrophilic (outer) region. (C) The result of docking simulation between eIF4A and 15d-PGJ2. The most plausible ligand binding site of eIF4A is highlighted inside the box. The hydrogen bonds between R295 of eIF4A and carboxyl tail of 15d-PGJ2 are presented in dotted red line.

By analyzing the docking simulation data of 15d-PGJ2-eIF4A, we also found that R295 residue of eIF4A might interact strongly with 15d-PGJ2 and makes the hydrogen bond (Fig. 2C). We found that our homology model is a reasonable surrogate for eIF4A structure since nearly all ligand interacting residues in the original complex structure are conserved in our model (Fig. 2D). Thus, we suggest that the hydrogen bond between the tail of 15d-PGJ2 and R295 residue of eIF4A might be susceptible to stabilize the flexible alpha-chain of 15d-PGJ2 and to aid the chain to dock easily with eIF4A. This simulation data suggests us that R295 can be an important target residue as 15d-PGJ2 recognizes eIF4A and binds to it.

Next, we asked to test whether the relationship between C264 and R295 is conserved through evolution. It is known that the residues that play structurally or functionally important roles within protein are evolutionary conserved and have high covariance values (Lockless and Ranganathan, 1999; Süel et al., 2003). To investigate the functional importance of C264 and R295, we calculated the covariance value for all residue pairs using homologues of human eIF4A1 (Fig.S2B) (see *Methods*). The histogram of cumulative counts shows that most pairs of residues have no strong correlations, however the covariance value of the C264-R295 pair is within the top 10% within eIF4a (Fig. S2A). This result suggests that both C264 and R295 participate together in an important biological function, that may include binding to 15d-PGJ2.

To experimentally confirm the structural relevance of the interaction between C264/R295 of eIF4A and 15d-PGJ2, we generated a C264S and R295A mutant of eIF4A. Binding of R295A mutant with 15d-PGJ2 is not reduced compared with wild type eIF4A, rather it increased slightly (Fig. 3A, lanes 1 and 3). However, binding of 15d-PGJ2 with C264S/R295A double mutant of eIF4A is significantly reduced compared with C264S mutant of eIF4A (Fig. 3A, lane 4), suggesting that R295 region has an additive function in stabilizing the interaction between 15d-PGJ2 and eIF4A.

**Figure 3.**
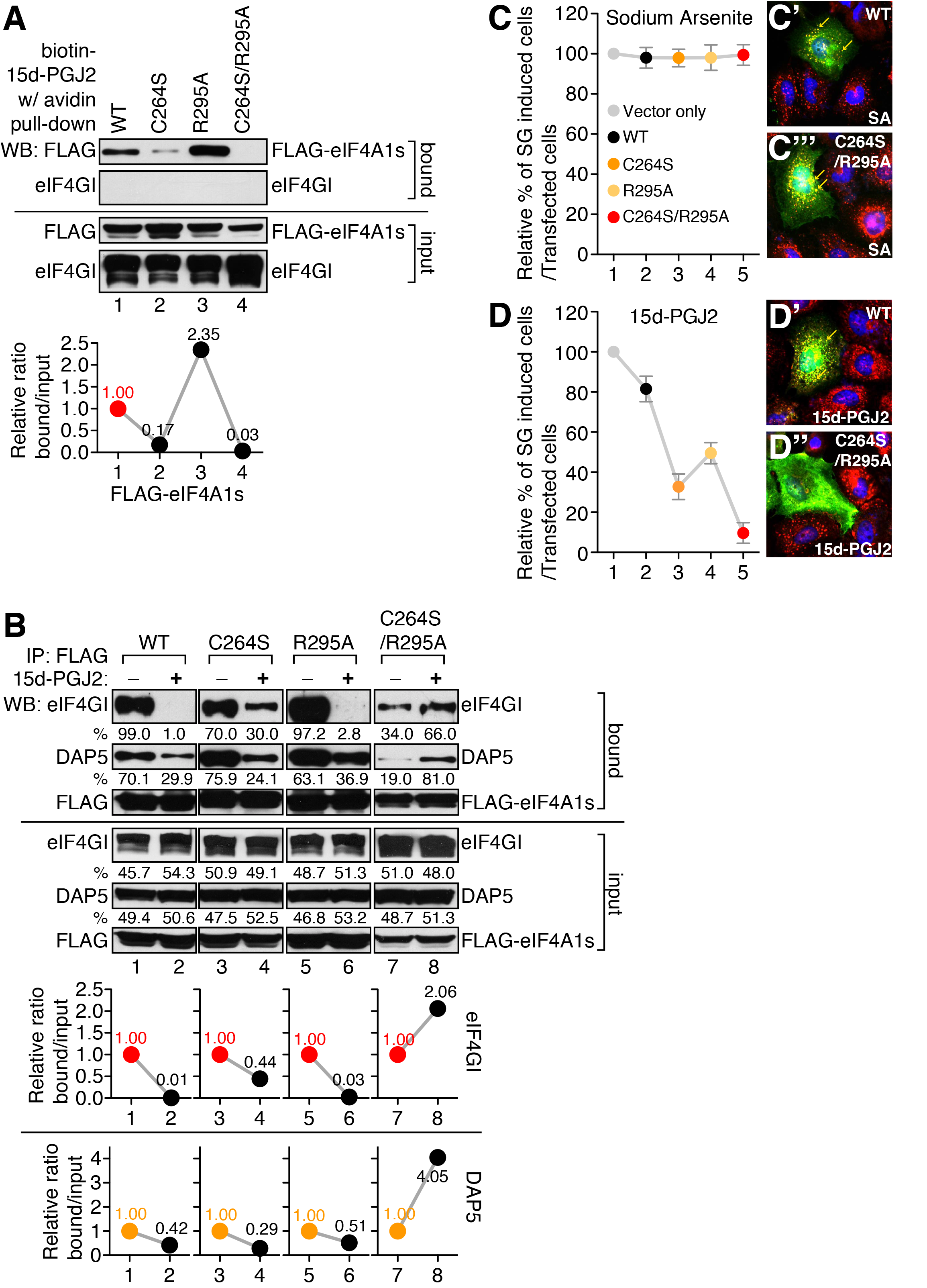
Binding of 15d-PGJ2 to arginine 295 of eIF4A is important for interaction with eIF4G and stress granules (SGs) formation. (A) 293T cells were transfected with the wild-type (WT, lane 1) or mutants (lanes 2-4) FLAG-eIF4A1s. Biotin pull-down assay was performed using biotin-15d-PGJ2 as described in **EXPERIMENTAL PROCDURES**. Western-blot analysis was performed with anti-FLAG and anti-eIF4GI antibodies. The bands of each lane are quantified using ImageJ as described in **EXPERIMENTAL PROCDURES**. Then the relative ratio of bound/input is calculated and visualized as graph below the band. (B) 293T cells were transfected with the wild-type (WT, lanes 1 and 2) or mutants (lanes 3 to 8) FLAG-eIF4A1. Immunoprecipitation was performed in the absence or presence of 15d-PGJ2 as described in **EXPERIMENTAL PROCDURES**. Western-blot analysis was performed with anti-FLAG, anti-DAP5, anti-eIF4GI antibodies. The bands of each lane are quantified using ImageJ as described in **EXPERIMENTAL PROCDURES**. Then the relative ratio of bound/input is calculated and visualized as graph below the band. (C) and (D) HeLa cells were grown on cover slips and transfected with a FLAG vector, wild type eIF4A (WT), or mutant eIF4As (C264S, R295A, and C264S/R295A). After 48 hours of incubation, cells were treated with the 400 μM of sodium arsenite (SA) (C) or 100 μM of 15d-PGJ2 (D) for 30 minutes. Cells were fixed and immunocytochemical analyses were performed with anti-FLAG and anti-eIF3b antibodies. SGs were counted among FLAG-eIF4As transfected cells. Each circle was normalized with vector transfectant. (C’), (C’’), (D’), and (D’’) Samples counted in pannel (C) and (D) were visualized. FLAG-eIF4As are green, eIF3B is red. The nuclei are shown in blue by Hoechst staining. SGs are marked as yellow arrows.

Next, we tested the effect of R295A mutation on the interactions between eIF4A and eIF4GI, which is inhibited by the binding of 15d-PGJ2 to eIF4A. The binding of C264S mutant eIF4A to eIF4GI was comparable to WT eIF4A (lane 3 of Fig 3B). However, the inhibitory effect of 15d-PGJ2 was dramatically reduced by that mutation (lanes 3 and 4 of Fig. 3B). The binding of R295A mutant eIF4A to eIF4GI was comparable to WT eIF4A (lane 5 of Fig. 3B), and the inhibitory effect of 15d-PGJ2 was also similar to WT (lanes 5 and 6 of Fig. 3B). When we tested the interaction between C264S/R295A double mutant eIF4A and eIF4GI, we found a slight decrease of the interaction (lane 7 of Fig. 3B). This interaction was not inhibited by 15d-PGJ2 treatment, suggesting that 15d-PGJ2 cannot bind to double mutant eIF4A and thus cannot block its interaction with eIF4G (lanes 7 and 8 of Fig. 3B). All these data suggest that R295 of eIF4A is an important target residue for 15d-PGJ2, which can regulate the interactions between eIF4A and eIF4GI.

To test the role of C264 and R295 residues of eIF4A in 15d-PGJ2-mediated physiological responses, we measured the numbers of SGs in transfected cells with different eIF4A constructs. We previously identified that the anti-inflammatory effect of 15d-PGJ2 partly results from inhibition of translational initiation (Kim et al., 2007) and SGs formation are good indicator of translational initiation blockage. We decided to compare the effect of sodium arsenite on SGs formation to that of 15d-PGJ2 since the SGs inducing-mechanisms of these two agents are distinct (Kim et al., 2007). SA induces SGs formation via phosphorylation of eIF2α, which inhibits efficient GDP-GTP exchange, leading to a decrease in levels of translationally competent eIF2/GTP/tRNA_i_^Met^ ternary complex, and inhibits translation initiation. In contrast to SA, 15d-PGJ2-dependent SGs formation is independent of eIF2α phosphorylation. Rather it targets eIF4A and inhibits the interaction between eIF4A-eIF4G, leading to inhibition of translation initiation (Panas et al., 2016).

We found that SA-induced SGs formation is not affected either by wild-type or double-mutant eIF4A overexpression (Fig. 3C), indicating that SA-dependent, in other words, eIF2α-dependent SGs formation is not affected by any forms of eIF4A overexpression. However, when the wild-type eIF4A was overexpressed, it could reduce the formation of SGs induced by 15d-PGJ2 up to 20% (lane 2 of Fig. 3D). When the mutant forms of eIF4As that do not bind to 15d-PGJ2 were overexpressed, 15d-PGJ2-dependent SGs formation was reduced by up to 90% (Fig. 3D). These data suggest that 15d-PGJ2-dependent SG formation is highly dependent on its binding to the C264 and R295 residues of eIF4A.

### C264 of eIF4A is highly conserved from worm to human

eIF4A and mechanisms of translation initiation are conserved across many species. If 15d2-PGJ2 regulation of eIF4A is important across species then its binding sites on eIF4A should be conserved. To define the evolutionary importance of these two amino acids in eIF4A, we compared the 15d-PGJ2 binding region of eIF4A among various species from budding yeast to human (Fig. S2B). In most species, C264 of eIF4A is highly conserved, however, in budding yeast *S. cerevisiae*, C264 is converted into tryptophan (Fig. S2B). In *D. melanogaster* and *C. elegans*, R295 is converted into histidine and asparagine, respectively (Fig. S2B). Thus, we suggest that C264 and R295 of eIF4A is relatively conserved through various species due to the 15d-PGJ2 actions on inflammation partly through translational blockage.

We next decided to examine the effect of 15d-PJ2 on various species showing the different amino acid pairs in the eIF4A region. To test the possible action of 15d-PGJ2 on translational blockage through eIF4A among various species, we chose several species and performed a series of experiments. First, we treated zebrafish embryos with 15d-PGJ2 at an early stage (4 hpf, hours after fertilization) and found that it results in gastrulation defects. We introduced two molecular markers; *chd*, an involuting dorsal mesoderm marker and *myod*, as an adaxial and somite marker, respectively (Fig. 4A, right panels). In addition, zebrafish embryos treated with 15d-PGJ2 at a later stage (10 hpf) caused a severe defect in spinal cord development at 28 hpf (Fig. 4A, red dashed lines in bottom panels).

**Figure 4.**
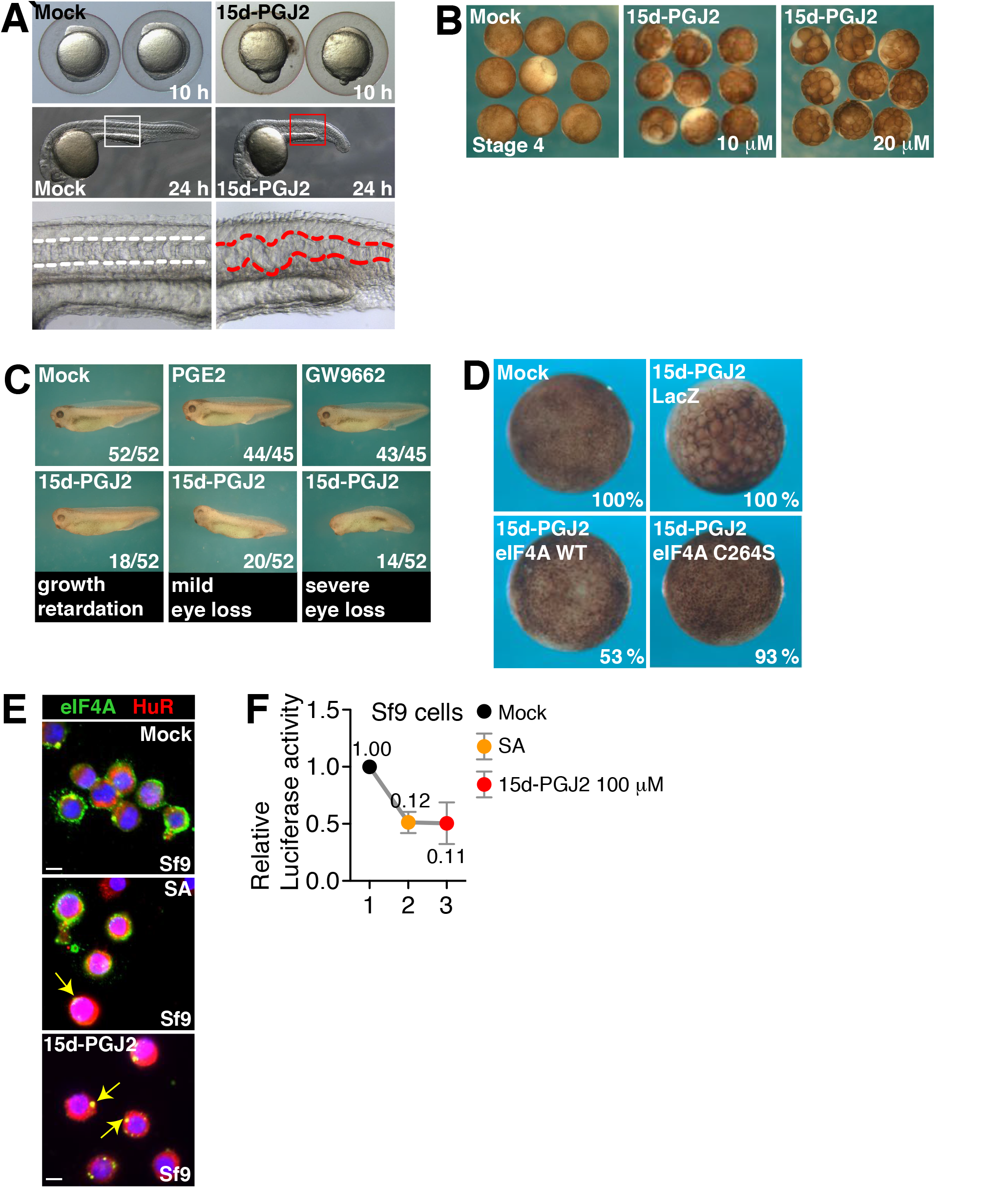
The effect of 15d-PGJ2 on various species. (A) Zebrafish embryos were mock treated or treated with 10 μM of 15d-PGJ2 at two different developmental stages (4 hpf or 10 hpf) and examined at later stages. 15d-PGJ2 treatment caused a gastrulation defect at early stage (6ss, 6 somite stage) and spinal cord defect at later stage (28 hpf), respectively. Early effects of 15d-PGJ2 were confirmed by using two molecular markers; *chd* (85% epiboly stage; L, lateral view; D, dorsal view) and *myod* (8ss, dorsal view). (B) *Xenopus* embryos were mock treated or treated with 20 μM of 15d-PGJ2 from stage 4 and cultured until gastrula stage (stage 11). Note that embryos treated with 15d-PGJ2 were developmentally arrested at early blastula stage. (C) *Xenopus* embryos were mock treated or treated with 20 μM of 15d-PGJ2, PGE2, or GW9662 after the onset of gastrulation (stage 11). Phenotypes were counted at stage 35. Among 52 embryos treated with 15d-PGJ2, 18 were growth retarded as shown by reduced trunk pigmentation and delayed eye formation, 20 showed mild eye defects, and 14 showed loss of eye and defects in dorsal axis. (D) *Xenopus* embryos were injected with β-galactosidase mRNA, eIF4A mRNA, or eIF4A C264S mutant mRNA at stage 2 and mock treated or treated with 20 μM of 15d-PGJ2 from 16-cell or 32-cell stages. Embryos were cultured until stage 11 and fixed. β-galactosidase mRNA injection or mock treatment was performed for the negative control. eIF4A injection rescued developmental arrest induced by 15d-PGJ2 administration (9/17, 53%), as well as did eIF4A C264S injections (14/15. 93%). (E) Sf9 cells were grown on cover slips and mock-treated (top panel), treated with 400 μM of SA (middle panel), or 50 μM of 15d-PGJ2 (bottom panel). Immunocytochemical analyses were performed with anti-eIF4A1 (green) and anti-HuR (red) antibodies. The nuclei are shown in blue by Hoechst staining. (F) Sf9 cells were co-transfected with monocistronic mRNAs containing renilla luciferase with cap-structure and firefly luciferase with CrPV IRES. After 4 h of trasnfection, cells were mock treated (lane 1), treated with 400 μM of SA (lane 2), or 100 μM of 15d-PGJ2 (lane 3) for 1 h. Luciferase assay was performed and relative luciferase activity was shown. Rluc/Fluc ratio means eIF4A dependent translation.

Second, to understand the function of eIF4A in 15d-PGJ2-mediated translational modulation, we chose *Xenopus* embryos to genetically manipulate the expression of eIF4A and its mutant forms. We found that Xenopus embryos also showed developmental defects when treated with 15d-PGJ2 (Fig. 4B). When 15d-PGJ2 was administered at a later developmental stage in *Xenopus*, most of the animals showed growth retardation, mild eye loss or severe eye loss (Fig. 4C, bottom panels and Fig. S4A) compared with mock-treated animals, or animals treated with a control prostaglandin (PGE2) or GW9662, potent PPAR-γ antagonist. Interestingly, developmental defects induced by 15d-PGJ2 were rescued by eIF4A mRNA injection (Fig. 4D). Inhibition of developmental defects by eIF4A mRNA injection might be resulted by buffering 15d-PGJ2 with overexpressed eIF4A (Fig. S4B). In addition, C264S mutant eIF4A mRNA injection almost completely rescued the developmental defect induced by 15d-PGJ2, suggesting that the binding motif of 15d-PGJ2 in eIF4A is critical for this developmental defect (Fig. 4D).

Third, we moved to test the effect of 15d-PGJ2 in invertebrate models. The eIF4A of fruit fly contains C264/H295 (Fig. S2B). We found that *Spodoptera frugiperda-* derived Sf9 cells form SGs-like structure by SA and 15d-PGJ2 using antibodies against eIF4A and RNA binding protein, HuR (Fig. 4E, yellow arrows on the right panel). By searching sequence database, we found that the 15d-PGJ2 binding sites of eIF4A in fall armyworm is conserved as C264/R295 form (Fig. S4C). To confirm the effect on the cap-dependent translation of SF9 cell by these chemicals, we transfected dual luciferase mRNA in Sf9 cell line then treated them with either SA or 15d-PGJ2. Interestingly, we found a strong correlation between luciferase assay and immunocytochemical data (Fig. 4F), indicating that 15d-PGJ2 can affect cap-dependent translation in insect cell. It has been reported that metazoan SGs assembly is mediated by P-eIF2α-dependent and –independent mechanisms (Farny et al., 2009), which is consistent with our data (Fig. 4E and F). The phosphorylation of eIF2α induces stress signals in Sf9 cells (Aarti et al., 2010), which is also consistent with our data (Fig. 4E and F). All these data and reports suggest that 15d-PGJ2 can induce SGs formation resulting in inhibited cap-dependent translation in invertebrate cells that contains conserved eIF4A amino acid residues for binding to 15d-PGJ2.

## DISCUSSION

Here we showed the specific effect of 15d-PGJ2 on eIF4A is dependent on conservation of C264 and amino acid residues in 295 position. First, we showed that binding of 15d-PGJ2 to eIF4A mostly blocks the interaction between MIF4G domain of eIF4G and eIF4A. We showed that the interactions between eIF4E-eIF4G, eIF3-eIF4G, eIF4G-PABP, and eIF4A-eIF4B were not affected by 15d-PGJ2 treatments (Fig. 1A, Fig. S1A-B). It is known that eIF4G contains two eIF4A binding motif, MIF4G and MA3 domain respectively. Using a domain mapping strategy, we showed that MIF4G domain is responsible for the inhibitory effect on the interactions between eIF4A and eIF4GI by 15d-PGJ2 (Fig. 1C-E, Fig. S1C-G). We also successfully showed that binding of 15d-PGJ2 to eIF4A inhibits the interaction between MIF4G domain and eIF4A by using various eIF4G homologues and eIF4A binding partners (Fig. 1F-I). Using a docking study, we identified that C264 of eIF4A can be accessible to 15d-PGJ2 (Fig. 2B) and found a clue that the tail of 15d-PGJ2 may locate very close to R295 of eIF4A (Fig. 2C). Although it is known that thiol modification of 15d-PGJ2 is enough for direct binding to its targets, we suggest that the tail of 15d-PGJ2 can stabilize or initiated the binding to targets. We next confirmed the predictions of our computer docking simulation with experiments. Our data suggest that R295 of eIF4A plays a critical role to stabilize the binding of 15d-PGJ2 and eIF4A (Fig. 3). To confirm that the conservation of C264/R295 plays an important role in translational regulation (Fig. S2A), we treated a variety of species with 15d-PGJ2 (Fig. S2B). We found interesting relations between C264/R295 conservation and 15d-PGJ2-mediated cell growth inhibition, translational inhibition, and/or SGs formation (Fig. 4 and Fig. S4). In summary, our study shows that 15d-PGJ2 specifically binds to C264/R295 residues of eIF4A and these binding properties are related to 15d-PGJ2-mediated translational inhibition.

Powerful computational simulation based docking studies are widely used to design and modify drugs. Using this technique, we could predict that the carboxyl tail of 15d-PGJ2 binds close to the R295 of eIF4A. As we predicted, previous reports by other groups also suggested that the carboxyl group of 15d-PGJ2 makes strong hydrogen-bonding interactions through lysine or arginine of target proteins (Pande and Ramos, 2005). Moreover, the carboxyl of PG-like fatty acids has been experimentally considered as an important determinant for molecular recognition with their natural receptors (Nolte et al., 1998; Xu et al., 1999). Previous study of eIF4A showed that D265 and D296 play a key role in its binding to eIF4G (Oberer et al., 2005). Crystal structure of the yeast eIF4A-eIF4G complex revealed that eIF4G-S612 makes a hydrogen bond to eIF4A-T252 (corresponding to C264 in human eIF4A (Oberer et al., 2005)) and eIF4G-E628 and D595 form a salt bridge with eIF4A-K284 (corresponding to D296 in human eIF4A (Oberer et al., 2005; Schütz et al., 2008). Whether the function of the 15d-PGJ2 tail is to affect stabilization or initiation of binding to eIF4A before covalent modification is not known, our experimental data shows that R295 of eIF4A is important for the binding of 15d-PGJ2 (Fig. 3). Since 15d-PGJ2 has multiple targets (Pande and Ramos, 2005), a docking study and experimental studies with these targets will be required to confirm the role of the 15d-PGJ2 tail region on interactions with other targets.

What is the implication of these molecular interactions on translational regulation? There are several interesting perspectives provided by the molecular details from our study. First, our data suggest that the binding of 15d-PGJ2 to eIF4A shows highly specific effects on cellular physiology. For example, it only affects translational initiation step thus inducing SGs formation. The binding of 15d-PGJ2 to eIF4A only blocks the interactions between eIF4G-eIF4A, not eIF4E-eIF4G, eIF3-eIF4G, or eIF4A-eIF4B (Fig. 1A, Fig. S1B). Rather, 15d-PGJ2 binding to eIF4A increases the interactions between eIF4A and RNA (Kim et al., 2007) thus resulting in the increases of the interactions between eIF4A and PABP (Fig. S1A), which accounts for the RNA-mediated interaction. Interestingly, the helicase activity of eIF4A was not altered by 15d-PGJ2 treatment (Fig. S3), indicating that the binding of 15d-PGJ2 to eIF4A specifically blocks its interactions with eIF4G.

It has been identified that MIF4G domain also called the HEAT-1 domain is mainly responsible to binding of eIF4G to the eIF4A C-terminus (237-406) (Marintchev et al., 2009). We also found that 15d-PGJ2 treatment only block the interactions between eIF4A and eIF4GI, PAIP1, and DAP5 but not the interactions between eIF4A-PDCD4 and eIF4A-eIF4GII. PDCD4 is a newly characterized tumor suppressor gene and functions by isolating eIF4A from eIF4F complexes (Yang et al., 2003). If 15d-PGJ2 only blocks translation through MIF4G domain but not the function of PDCD4, the anti-proliferative effect of 15d-PGJ2 and PDCD4 may be cumulative within cells. The interactions between eIF4A-eIF4GII was relatively weak and not affected by 15d-PGJ2 treatment (Fig. S1C-G). eIF4GI and eIF4GII have differing functions as reported by other groups (Svitkin et al., 1999), however, we are unsure why 15dPGJ2 may not bind eIF4GII. Further study is required to find the molecular details of this phenomenon.

Cyclopentenone prostaglandins are produced at the late stage of inflammatory responses to stop the positive feedback loop and prevent sustained inflammation (Straus et al., 2000; Straus and Glass, 2001). Previously, we suggested that the antiinflammatory action of 15d-PGJ2 partially resulted from translational inhibition. We also suggested that eIF4A is a possible candidate for that function of 15d-PGJ2 (Kim et al., 2007). If this were the case, C264 of eIF4A would be critical for 15d-PGJ2 actions. Evolutionary conservation of the C264 of eIF4A among mammals and many multicellular organisms motivated us to test this possibility (Fig. S2B). From mammals to insects, 15d-PGJ2 inhibited cap-dependent translation (Fig. 4A), induced SGs like structures (Fig. 4E), or disrupted development (Fig. 4A-D). We cannot conclude all these phenomena result from 15d-PGJ2 effects on eIF4A; however, 15d-PGJ2 treatment of *Xenopus* embryos with eIF4A mRNA could prevent developmental defects induced by 15d-PGJ2 (Fig. 4D). We also confirmed that a C264S mutant eIF4A, that is resistant to 15dPGJ2, can rescue the developmental delay better than WT eIF4A, suggesting that binding specificity of 15d-PGJ2 to C264 of eIF4A is critical for developmental delays in *Xenopus* embryo. Thus, overexpression of eIF4A can rescue the developmental defects induced by 15d-PGJ2 at least in *Xenopus* embryos.

15d-PGJ2 is synthesized by dehydration of PGD2. PGD2 synthesis requires PGD2 synthases: HPGDS (entrezID: 27306) and LPGDS (entrezID: 5730) (Scher and Pillinger, 2005). The existence of PGD2 synthases in the genome of species could be a possible criteria of 15d-PGJ2 production in that species. To test this possibility, we searched for the orthologues of human PGDS using InParanoid (Fig. S5). The orthologues of PGDS are found in mouse, *Xenopus* and *Drosophila* in which C264 is conserved. In those species, 15d-PGJ2 can induce SG-like structures (Fig. 4 E), disrupt development (Fig. A-D), or inhibit translation (Fig. 4F). In zebrafish, however, orthologues of PGDS were not found though the treatment of 15d-PGJ2 induces developmental defects (Fig. 4A). This suggests that the possible existence of different enzymes producing PGD2 in zebrafish. Although there is the exceptional case such as zebrafish, the effect of 15d-PGJ2 seems to be correlated with the existence of PGDS in the genome of the species.

Our finding can provide the strategy to design more efficient drug. For example, covalent modification of HIV Tat protein by 15d-PG2 can be applied to design anti-viral drugs. Since 15d-PGJ2 has specific cellular target proteins, finding targets and designing more efficient structures will be helpful for medical applications such as Ischemia reperfusion (Blanco et al., 2005; DeGracia et al., 2006; Kayali et al., 2005; Kloner and Rezkalla, 2006; Lin et al., 2006; McDunn and Cobb, 2005; Murry et al., 1986). Together with small molecules showing anti-cancer effects such as Pateamine A and 4EGI-1 (Korneeva et al., 2001), we suggest that targeting the process of translational initiation could be a reasonable strategy to improve anti-cancer and anti-inflammatory treatments.

## EXPERIMENTAL PROCEDURE

### Plasmid construction

Plasmid information is described elsewhere (Kim et al., 2007) (Kim et al., 2005). Site-directed mutagenesis was performed by *DpnI* selection method using proper primers. All plasmids are sequenced to confirm the mutagenesis.

### Antibodies and chemicals

Antibody against FLAG was purchased from Sigma, GFP and HA from Santa Cruz. Antibody against eIF4GI was prepared in our laboratory (Kim et al., 2005). Chemicals 15d-PGJ2, biotinylated 15d-PGJ2, and PGE2 were purchased from Cayman Chemical. Sodium arsenite was purchased from Sigma. Immobilized streptavidin agarose was purchased from Pierce.

### Quantification of Western blot analysis

We quantified the density of bands using ImageJ (http://rsb.info.nih.gov/ij/index.html) software. We created digital images of gels using digital scanner then follow the protocol for Gel analysis menu in ImageJ (Schneider et al., 2012). In short, we convert gel images to 8-bit images then choose the Rectangular Selections tool to draw a rectangle around the first each lane. After drawing the rectangles, we plot lanes using Plot Lanes menu then choose the peak using Straight Line selection tool. When all the peaks have been highlighted, we label peaks to express the percentage of each peaks compared to the total size of all the highlighted peaks. The quantification method using above method are described elsewhere (Gassmann et al., 2009; Tan and Ng, 2008).

### Cell cultures and transient transfection

HeLa cells and 293T cells were grown as described elsewhere (Kim et al., 2005).

### Pull-down with streptavidin and Immunoprecipitation

Biotin pull-down and immunoprecipitation experiments were performed as described elsewhere (Kim et al., 2007). In short, 293T cells transfected with DNAs were lysed using the NP-40 lysis buffer. The lysates were clarified by centrifugation at 14 000 g for 15 min. Anti-FLAG monoclonal antibody (4 μg) was incubated with 20 μl of Protein G agarose for 1 h in 1 ml NP-40 lysis buffer at 4°C. Lysates were pre-cleared with 10 μl of protein G agarose at 4°C for 30 min. After pre-clearing, cell lysates were treated with 50 μM of EtOH, PGE2, or 15d-PGJ2 at 30°C for 1 h, followed by centrifugation. Then protein G agarose☐conjugated antibodies were incubated with the pre-cleared lysates at 4°C for 1 h. Precipitates were washed three times with lysis buffer and analyzed by SDS–PAGE.

### Fluorescence microscopy

The immunocytochemical analyses of proteins were performed as described elsewhere (Kim et al., 2005). In short, after transfection of DNAs, cells were grown on coverslips coated with 0.2% gelatin for 48 h and then washed three times with phosphate-buffered saline (PBS). The cells were fixed with 3.5% (wt/vol) paraformaldehyde (Sigma) at room temperature (RT) for 12 min. After being washed three times with PBS, the cells were permeabilized with 0.1% Triton X-100 at RT for 2 min and then washed three times with PBS. The samples were soaked in blocking solution (PBS containing 1% bovine serum albumin) for 30 min at RT and then incubated with primary antibodies for 1 h at RT. After being washed with PBS, the samples were treated with Hoechst 33258 for 2 min at RT and washed again with PBS three times. Samples were treated with rhodamine tetramethyl isocyanate-conjugated and/or fluorescein isothiocyanate-conjugated secondary antibodies (Jackson ImmunoResearch Laboratories, Inc.) for 1 h at RT. Finally, the coverslips were washed three times with PBS, placed on a glass slide, and then sealed with transparent nail polish. The fluorescent images were captured with a cooled charge-coupled device camera and a Zeiss (Jena, Germany) Axioplan microscope. Data were processed using Image J software.

### Luciferase assay

Luciferase assays were performed as described elsewhere (Kim et al., 2005). In short, monocistronic mRNAs containing renilla luciferase with cap-structure and firefly luciferase with CrPV IRES was transfected into Sf9 cells by lipopectamine together. After 12 hrs of transfection, SA or 15d-PGJ2 was treated to cells for 1 h then luciferase assays were performed with a dual luciferase assay kit (Promega) per the manufacturer’s instructions. *Renilla* luciferase activity values were normalized by Firefly luciferase activity values that reflect transfection efficiency and general cellular activities.

### Insect cell cultures

BTI-TN-5B1-4 cells (High Five; Invitrogen) were maintained and transfected as described elsewhere (Farrell and Iatrou, 2004). Monocistronic Rluc and Fluc plasmids are described elsewhere (Kim et al., 2007).

### *Xenopus* embryo manipulation

*Xenopus* eggs were obtained and fertilized as described elsewhere (Kim and Han, 2007). Nieuwkoop and Faber stages were considered for the *Xenopus* developmental staging (Nieuwkoop and Faber, 1956). *In vitro* synthesized eIF4A mRNA was introduced into the *Xenopus* embryos by microinjection using Nanoliter injector (WPI). Embryos were cultured in 0.33X-modified ringer (MR) and treated with 20 μM of 15d-PGJ2 or GW9662 from the indicated stages.

### Zebrafish Experiment

Zebrafish were maintained at 28.5°C in a 14 h light/10 h dark cycle. Embryonic stages were determined by the hours post-fertilization (hpf) and microscopic observation. Zebrafish embryos were treated with 10 μM of 15d-PGJ2 at two different developmental stages; 4 hpf (before gastrulation) and 10 hpf (after gastrulation), respectively. Whole-mount *in situ* hybridization was performed as previously described (Jung et al., 2010).

### Homology modeling and docking

Crystal structure (1HV8) of MjDEAD extracted from the *hyperthermophile Methanococccus jannaschii* is used as a template to build the homology model of the human eIF4A1 (Oberer et al., 2005) using Modeller8v (Sali and Blundell, 1994). MjDEAD and human eIF4A1 shares high sequence identity and similarity, 33.8% and 54.4 % respectively. The sequences alignment between human eIF4A-1 and MjDEAD was performed using ClustalW (Chenna et al., 2003) and the alignment score were calculated from EMBOSS-align (Rice et al., 2000). Cα rmsd was calculated using MOE (ChemicalComputingGroup, 2008). The model structure of eIF4A-1 was energy-minimized using AMBER9 (Case et al., 2006). Since CYS is oxidized in the experimental environment, CYM potential parameters were used for CYS in the energy-minimized. We docked 15d-PGJ_2_ into the human eIF4A-1 model structure using eHits (Kerwin, 2005). eHits considers the flexibility of ligand and generates all possible ligand conformations, which has proven to be effective for modeling a ligand docking model. Before docking, 15d-PGJ_2_ was energy-minimized by *ab initio* quantum chemical calculation using Gaussian program (Frisch et al., 2004). The energy minimized 15d-PGJ_2_ was docked to the eIF4A-1 model.

### Calculation of covariance

We found 500 homologues of the human eIF4A-1 using wu-BLAST and filtered out 197 sequences whose length is smaller than 0.7 times or larger than 1.4 times of human eIF4A sequence and whose identity is greater than 90%. We removed all columns with gaps more than 50%. Finally, 303 selected sequences were aligned with human eIF4A sequences and the co-variance was calculated using ELS (Dekker et al., 2004) (Fodor and Aldrich, 2004).

### Conserved C-R pair in vertebrate orthologues of human eIF4A-1

We aligned 11 orthologues of human eIF4A-1 from InParanoid, Eukaryotic Ortholog Groups (Remm et al., 2001), using ClustalW.

### Searching Orthologues of PGDS

We search InParanoid, the Eukaryotic Ortholog Groups, for the orthologues of human PGDS (Remm et al., 2001), excluding inparalogues scoring below 0.05.

### Helicase assay

We performed *in vitro* helicase assay using 32P-labeld oligonucleotides as described elsewhere (Kim and Seo, 2009). The oligonucleotide sequences we used are described below. R-28-5’; 28mer: aaaacaaaacaaaauagcaccguaaagc and R-13; 13mer: gcuuuacggugcu.

## ACKNOWLEDGEMENTS

We are grateful to the members of Pohang Biology Group (PBG) for generous provision of advice and discussion. We also appreciate Dr. Derrick Gibbings (University of Ottawa) for his reading and reviewing this manuscript carefully. We also would like to thank Dr. Hyeshik Jang (Seoul National University) for finding the sequences of eIF4A from the *Spodoptera frugiperda* sequencing database. This work was supported by NRF (National Honor Scientist Program: 2010-0020414, WCU: R32-2008-000-10180-0) and KISTI (KSC-2011-G3-02).

---

**Supplement Figure 1.**
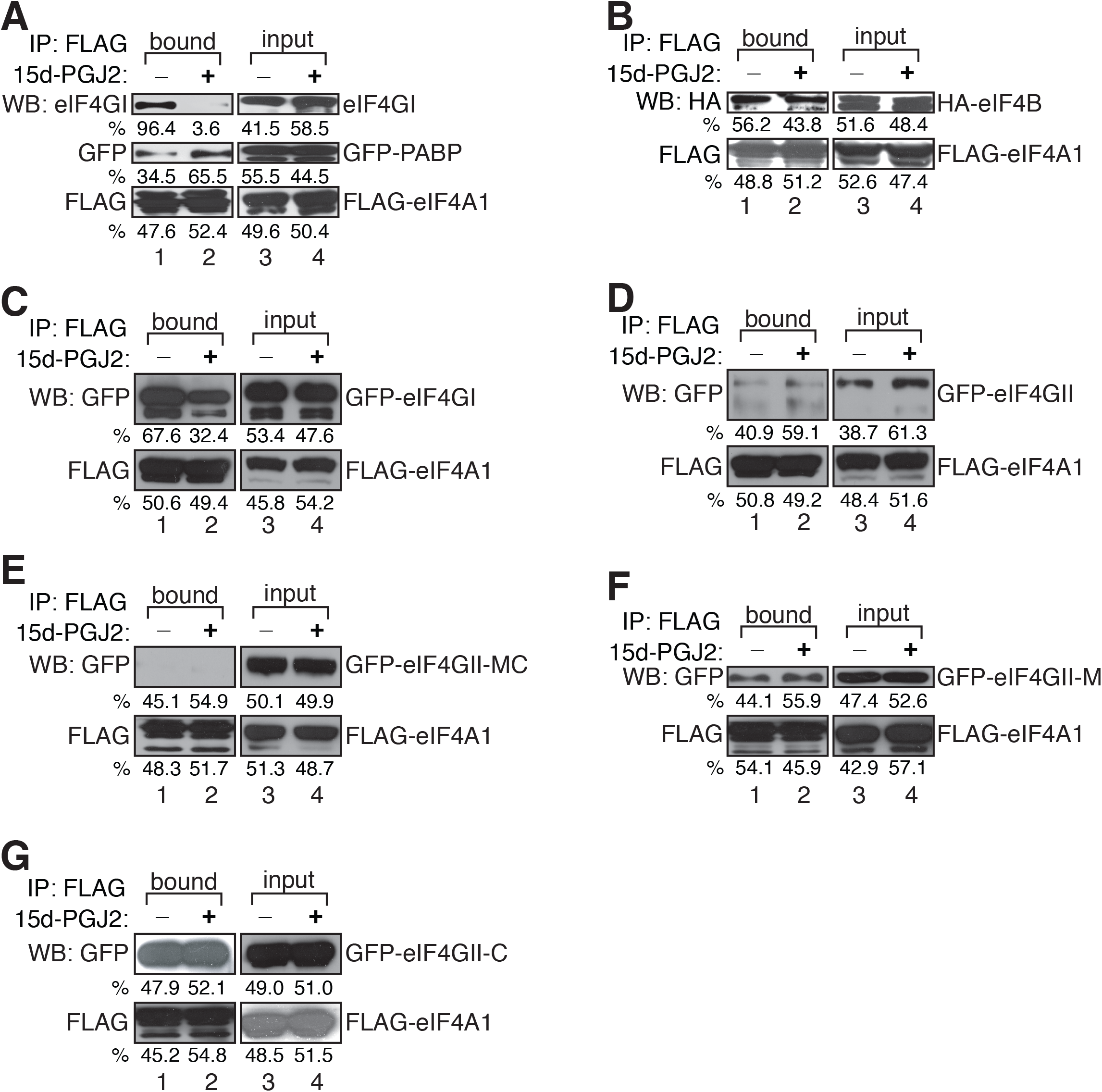
The effect of 15d-PGJ2 treatments on the interactions between translation initiation factors. (A) and (B) We have adopted this data from previous publication and reanalyzed it using ImageJ for quantification (Kim et al., 2007). (C)-(G) 293T cells were co-transfected with GFP-eIF4GI full length (C), GFP-eIF4GII full length (D), GFP-eIF4GII-MC (E), GFP-eIF4GII-M (F), and GFP-eIF4GII-C (G) and FLAG-eIF4A1. Cells were lysed then treated with EtOH or 10 μM of 15d-PGJ2 at 30°C for 1 hour. Immunoprecipitation was performed as described above then Western-blot analysis was performed with anti-FLAG and anti-GFP antibodies.

**Supporting Figure 2.**
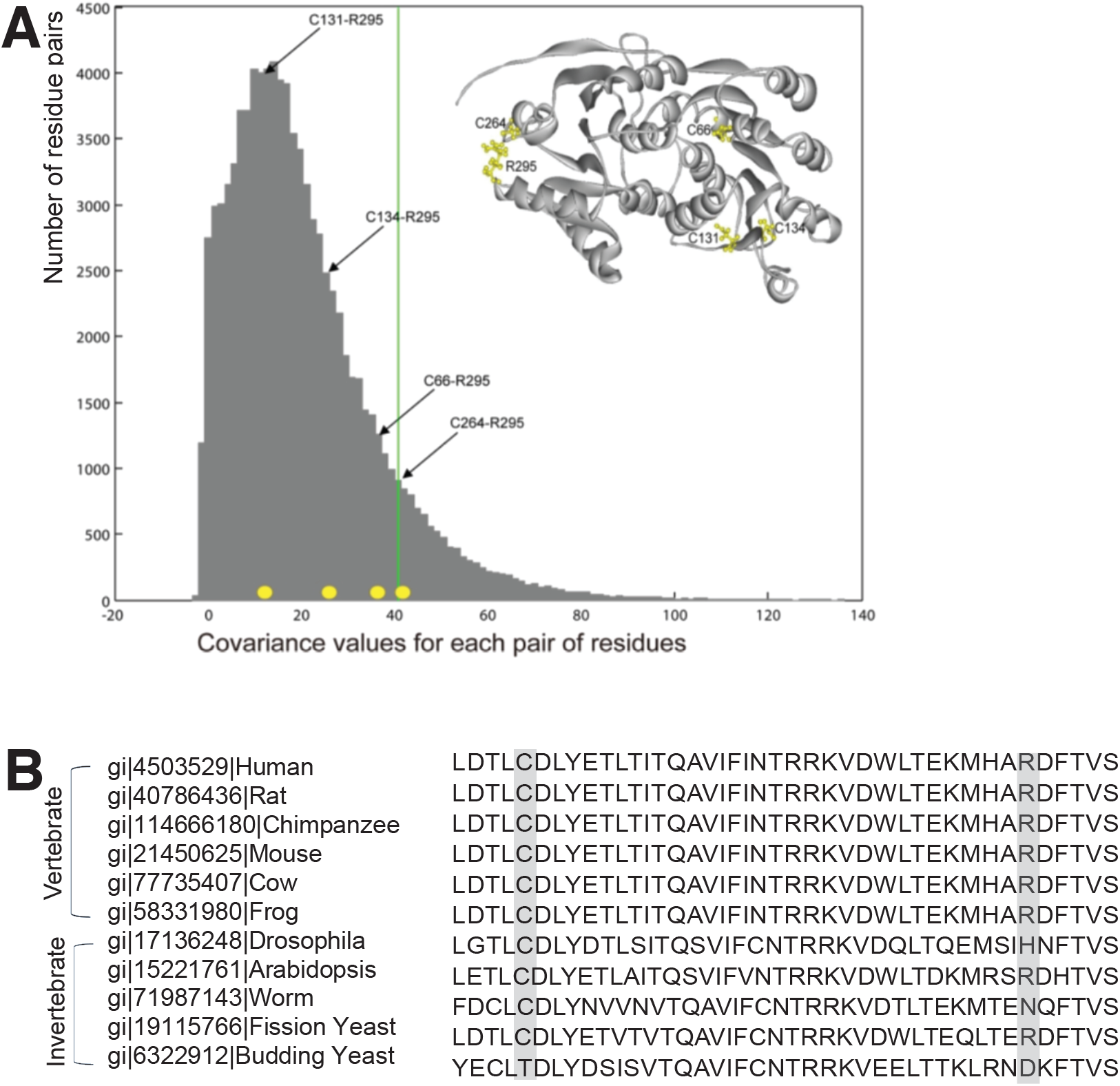
Covariance and conservation of C-R pair. (A) We used the eIF4AI structure to predict the solvent accessible surface. C264 was highlighted in yellow circle. (B) Covariance value for each pair of amino acid residues within eIF4A. The histogram shows the number of residue pairs corresponding to each covariance value. The green line represents the covariance value of top 10 percentile. The covariance values for C-R pairs are highlighted with yellow dots. The four C-R pairs are presented in the bracket. In the figure inside, four cysteins (C66, C131, C134, C264) and R295 are colored in yellow. (B) Conserved C-R pair in vertebrate orthologues of human eIF4A-1. The 11 orthologues of human eIF4A-1 from vertebrate and invertebrate are aligned. Cys and Arg are boxed in gray in alignment of 11 orthologues of human eIF4A-1. (D) The examples of the docking simulation results. The rank1 represents C264.

**Supplement Figure 3.**
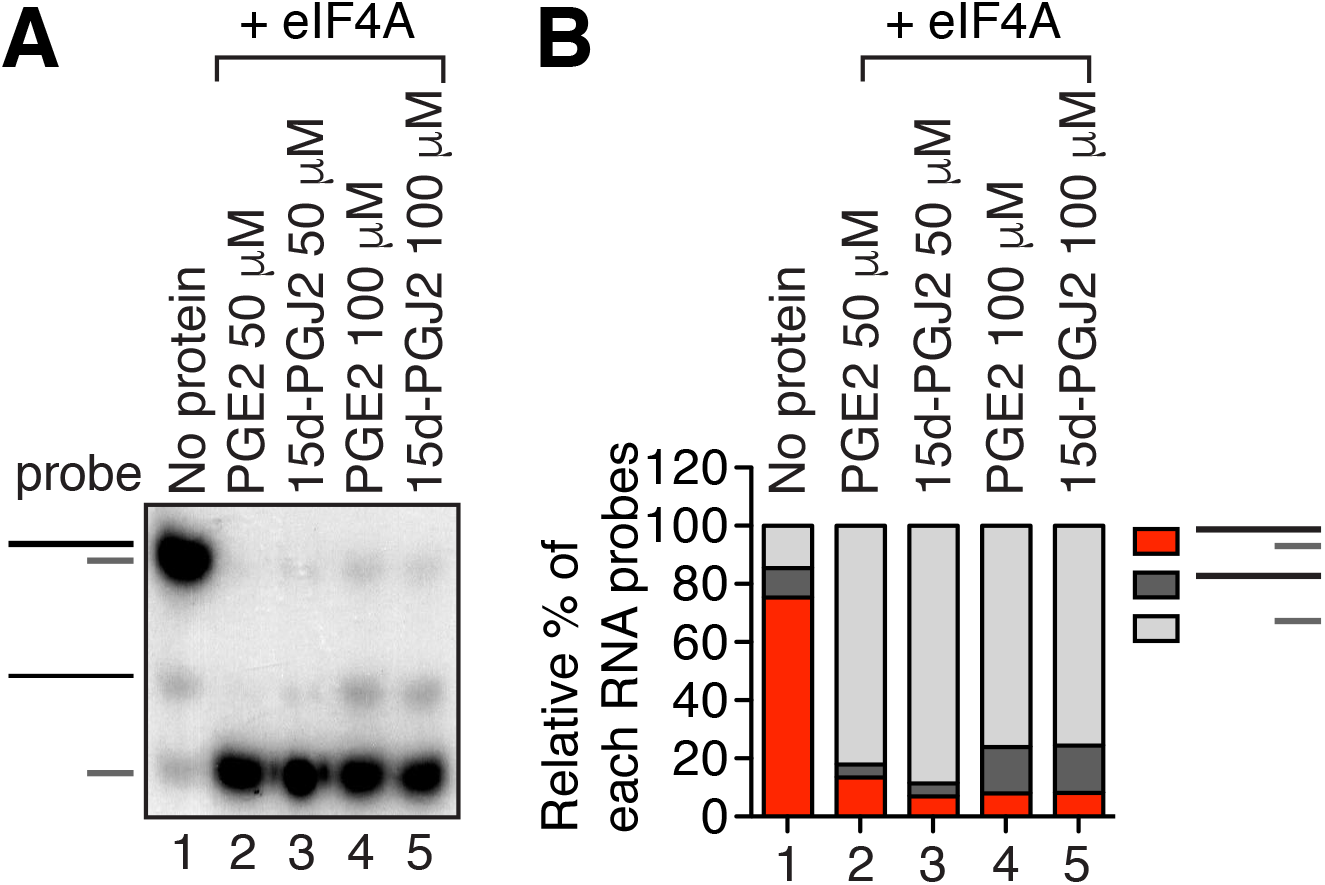
The effect of C264/R295 mutation on the 15d-PGJ2 binding to eIF4A. (A) Helicase assay was performed using purified His-eIF4A and radiolabeled oligonucleotides in the presence of PGE2 or 15d-PGJ2. (B) The gel images of (A) were analyzed with ImageJ and the relative amount of double strand, longer primer, and shorter primer was calculated.

**Supporting Figure 4.**
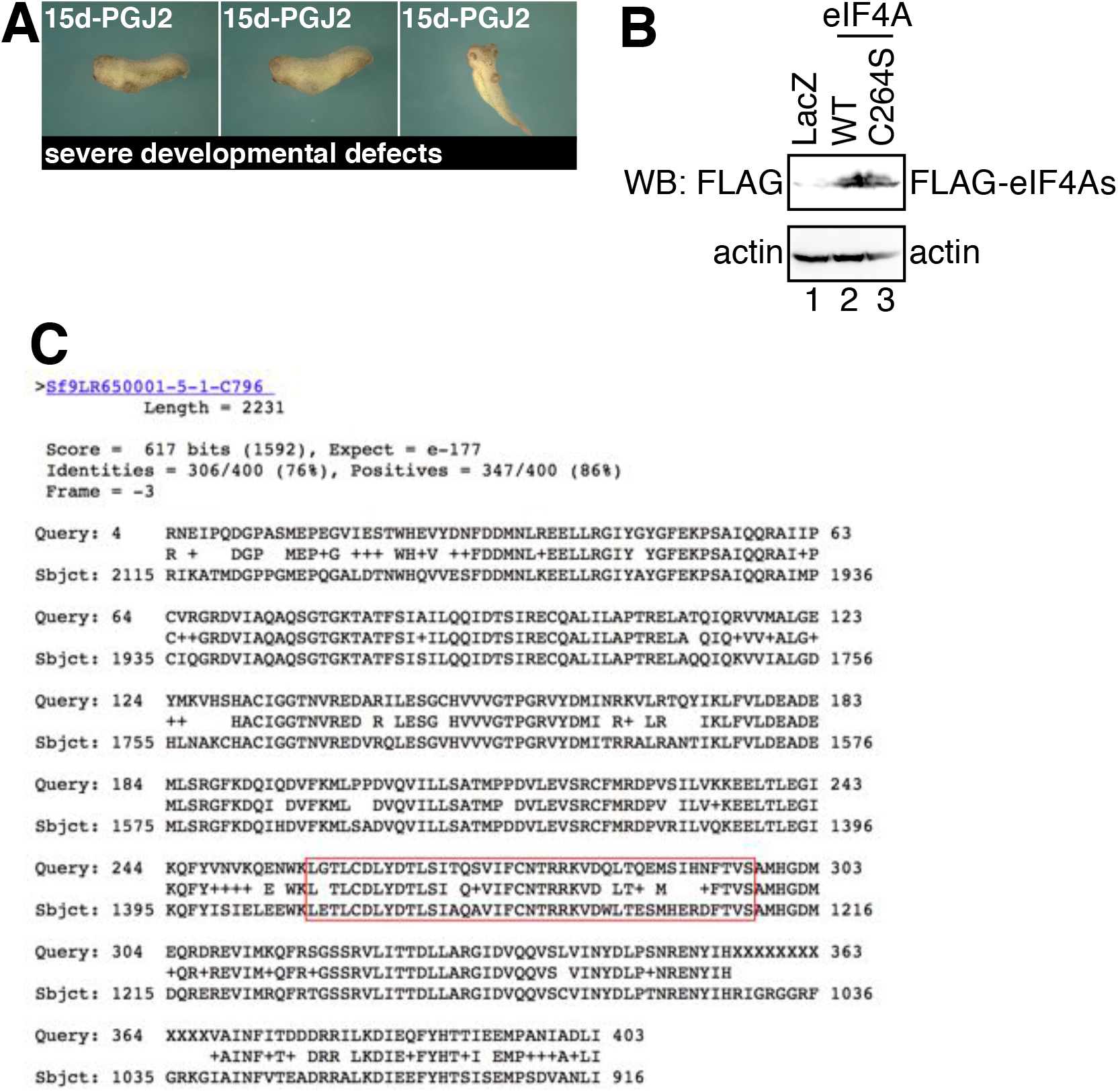
Continued data of the effect of 15d-PGJ2 on various species. (A) Examples of severe developmental defects of *Xenopus* by 15d-PGJ2 treatment. (B) The expression of FLAG-eIF4A mRNA used in Fig. 4E experiments were confirmed by western blot analysis. (C) Amino acid sequence of *Spodoptera frugiperda* eIF4A. Reference sequence was human eIF4A and the region contains C264~R295 is highlighted.

**Supplement Figure 5.**
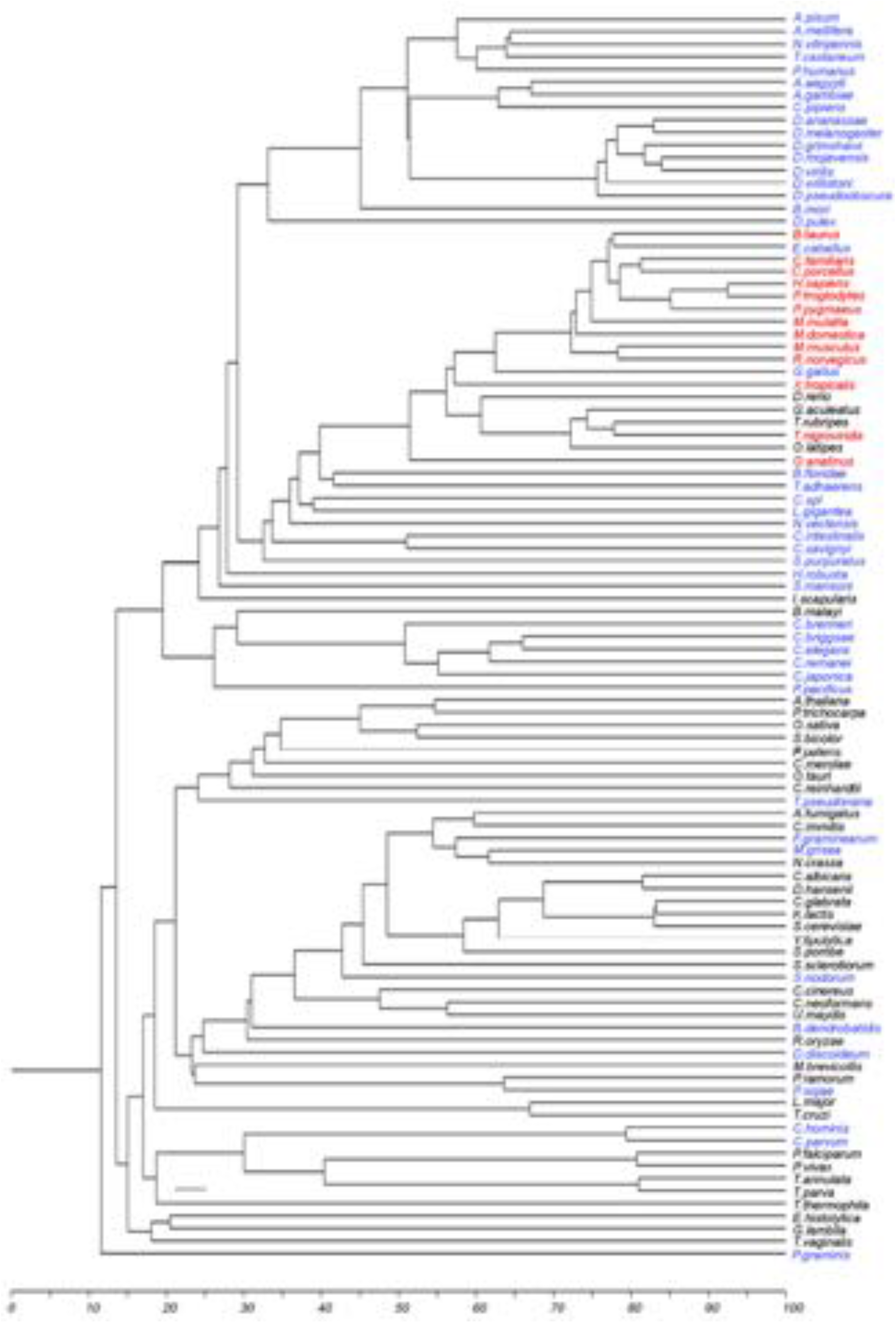
Orthophylogram of PGDS based on the average fraction of InParanoid orthologues between species (Remm et al., 2001). Blue/Red and black represent the species with or without the orthologues of human PGDS, respectively. Red represents the species that the orthologues of both PGDSs, HPGDS (entrezID: 27306) and LPGDS (entrezID: 5730), are found. Blue represents the species that the orthologues of HPGDS (entrezID: 27306) are found. Black represents the species that had no orthologues of PGDSs. (B) The sequence alignment of human eIF4A-1 with MjDEAD. The conserved motifs of the DEAD box helicase are highlighted with gray boxes.

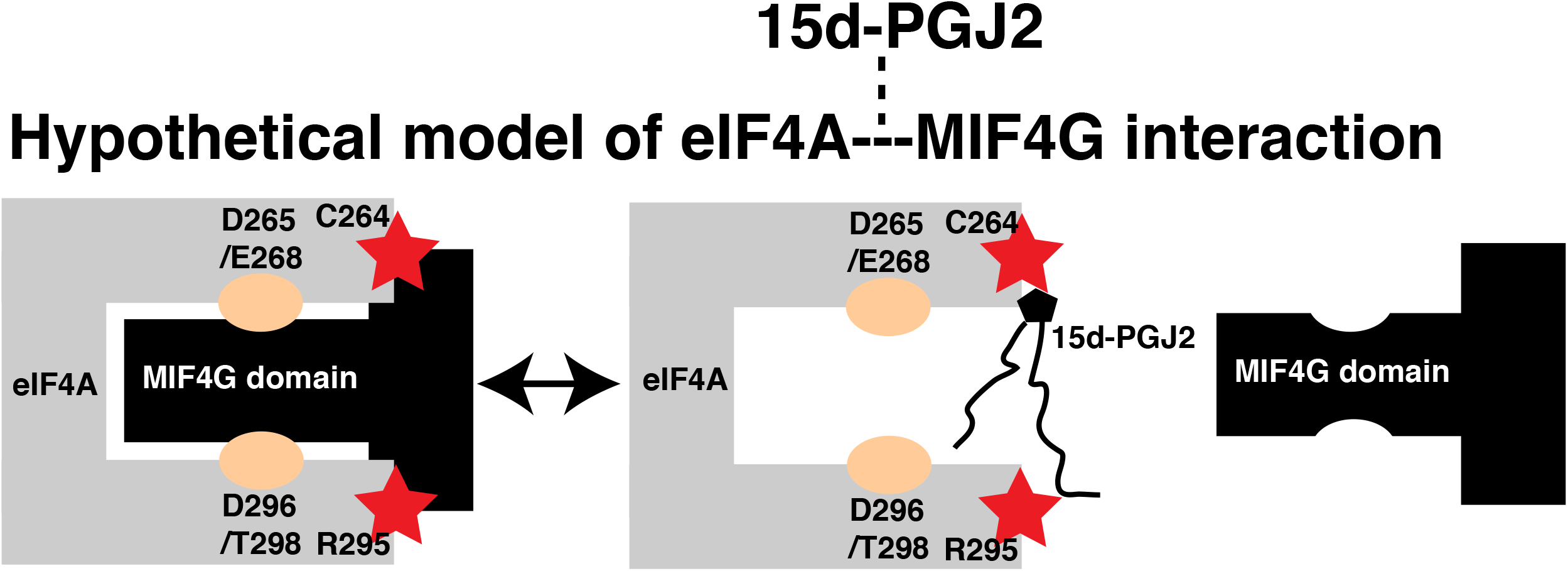

## REFERENCES

Aarti, I., Rajesh, K., Ramaiah, K.V.A., 2010. Phosphorylation of eIF2 alpha in Sf9 cells: a stress, survival and suicidal signal. Apoptosis 15, 679–692.

Blanco, M., Moro, M.Á., Dávalos, A., Leira, R., Castellanos, M., Serena, J., Vivancos, J., Rodríguez-Yáñez, M., Lizasoain, I., Castillo, J., 2005. Increased plasma levels of 15-deoxy Δ prostaglandin J2 are associated with good outcome in acute atherothrombotic ischemic stroke. Stroke 36, 1189–1194.

Case, D.A., Darden, T.A., Cheatham III, T.E., Simmerling, C.L., Wang, J., Duke, R.E., Luo, R., Merz, K.M., Pearlman, D.A., Crowley, M., 2006. AMBER 9. Univ. California, San Fr. 45.

ChemicalComputingGroup, M.O.E., 2008. Molecular Operating Environment.

Chenna, R., Sugawara, H., Koike, T., Lopez, R., Gibson, T.J., Higgins, D.G., Thompson, J.D., 2003. Multiple sequence alignment with the Clustal series of programs. Nucleic Acids Res. 31, 3497–3500.

Craig, A.W.B., Haghighat, A., Annie, T.K., Sonenberg, N., 1998. Interaction of polyadenylate-binding protein with the eIF4G homologue PAIP enhances translation. Nature 392, 520–523.

DeGracia, D.J., Rafols, J.A., Morley, S.J., Kayali, F., 2006. Immunohistochemical mapping of total and phosphorylated eukaryotic initiation factor 4G in rat hippocampus following global brain ischemia and reperfusion. Neuroscience 139, 1235–1248.

Dekker, J.P., Fodor, A., Aldrich, R.W., Yellen, G., 2004. A perturbation-based method for calculating explicit likelihood of evolutionary co-variance in multiple sequence alignments. Bioinformatics 20, 1565–1572.

Diaz-Camino, C., Risseeuw, E.P., Liu, E., Crosby, W.L., 2003. A high-throughput system for two-hybrid screening based on growth curve analysis in microtiter plates. Anal. Biochem. 316, 171–174.

Farny, N.G., Kedersha, N.L., Silver, P.A., 2009. Metazoan stress granule assembly is mediated by P-eIF2α-dependent and-independent mechanisms. Rna 15, 1814– 1821.

Farrell, P., Iatrou, K., 2004. Transfected insect cells in suspension culture rapidly yield moderate quantities of recombinant proteins in protein-free culture medium. Protein Expr. Purif. 36, 177–185.

Fodor, A.A., Aldrich, R.W., 2004. Influence of conservation on calculations of amino acid covariance in multiple sequence alignments. Proteins Struct. Funct. Bioinforma. 56, 211–221.

Frisch, Mj., Trucks, G.W., Schlegel, Hb., Scuseria, G.E., Robb, M.A., Cheeseman, J.R., Montgomery Jr, J.A., Vreven, T., Kudin, K.N., Burant, Jc., 2004. Gaussian 03, revision c. 02; Gaussian. Inc., Wallingford, CT 4.

Gassmann, M., Grenacher, B., Rohde, B., Vogel, J., 2009. Quantifying Western blots: pitfalls of densitometry. Electrophoresis 30, 1845–1855. doi:10.1002/elps.200800720

Imataka, H., Sonenberg, N., 1997. Human eukaryotic translation initiation factor 4G (eIF4G) possesses two separate and independent binding sites for eIF4A. Mol. Cell. Biol. 17, 6940–6947.

Jung, S., Kim, S., Chung, A., Kim, H., So, J., Ryu, J., Park, H., Kim, C., 2010. Visualization of myelination in GFP-transgenic zebrafish. Dev. Dyn. 239, 592–597.

Kayali, F., Montie, H.L., Rafols, J. a, DeGracia, D.J., 2005. Prolonged translation arrest in reperfused hippocampal cornu Ammonis 1 is mediated by stress granules. Neuroscience 134, 1223–45. doi:10.1016/j.neuroscience.2005.05.047

Kerwin, S.M., 2005. eHiTS 5.1. 6 SimBioSys Inc., 135 Queen’s Plate Drive, Unit 420, Toronto, Ontario M9W 6V1, Canada. http://simbiosys.ca/index.html. For pricing information, contact company. J. Am. Chem. Soc. 127, 8899–8900.

Kim, G., Han, J., 2007. Essential role for β-arrestin 2 in the regulation of Xenopus convergent extension movements. EMBO J. 26, 2513–2526.

Kim, J.-H., Seo, Y.-S., 2009. In vitro assays for studying helicase activities. Methods Mol. Biol. 521, 361–379. doi:10.1007/978-1-60327-815-7_20

Kim, W.J., Kim, J.H., Jang, S.K., 2007. Anti-inflammatory lipid mediator 15d-PGJ2 inhibits translation through inactivation of eIF4A. EMBO J. 26, 5020–5032.

Kim, W.J.W., Back, S.H.S., Kim, V., Ryu, I., Jang, S.K., 2005. Sequestration of TRAF2 into Stress Granules Interrupts Tumor Necrosis Factor Signaling under Stress Conditions. Mol. Cell. … 25, 2450–62. doi:10.1128/MCB.25.6.2450

Kloner, R.A., Rezkalla, S.H., 2006. Preconditioning, postconditioning and their application to clinical cardiology. Cardiovasc. Res. 70, 297–307.

Kondo, M., Shibata, T., Kumagai, T., Osawa, T., Shibata, N., Kobayashi, M., Sasaki, S., Iwata, M., Noguchi, N., Uchida, K., 2002. electrophile that induces neuronal apoptosis 2.

Koo, B., Lim, H., Chang, H.J., Yoon, M., Choi, Y., Kong, M., Kim, C., Kim, J., Park, J., Kong, Y., 2009. Notch signaling promotes the generation of EphrinB1-positive intestinal epithelial cells. Gastroenterology 137, 145–155.

Korneeva, N.L., Lamphear, B.J., Hennigan, F.L.C., Merrick, W.C., Rhoads, R.E., 2001. Characterization of the two eIF4A-binding sites on human eIF4G-1. J. Biol. Chem. 276, 2872–2879.

Lin, J.-C., Hsu, M., Tarn, W.-Y., 2007. Cell stress modulates the function of splicing regulatory protein RBM4 in translation control. Proc. Natl. Acad. Sci. 104, 2235– 2240.

Lin, T.-N., Cheung, W.-M., Wu, J.-S., Chen, J.-J., Lin, H., Chen, J.-J., Liou, J.-Y., Shyue, S.-K., Wu, K.K., 2006. 15d-prostaglandin J2 protects brain from ischemia-reperfusion injury. Arterioscler. Thromb. Vasc. Biol. 26, 481–487.

Lockless, S.W., Ranganathan, R., 1999. Evolutionarily conserved pathways of energetic connectivity in protein families. Science (80-. ). 286, 295–299.

Lomakin, I.B., Hellen, C.U.T., Pestova, T. V, 2000. Physical association of eukaryotic initiation factor 4G (eIF4G) with eIF4A strongly enhances binding of eIF4G to the internal ribosomal entry site of encephalomyocarditis virus and is required for internal initiation of translation. Mol. Cell. Biol. 20, 6019–6029.

Low, W.-K., Dang, Y., Schneider-Poetsch, T., Shi, Z., Choi, N.S., Merrick, W.C., Romo, D., Liu, J.O., 2005. Inhibition of eukaryotic translation initiation by the marine natural product pateamine A. Mol. Cell 20, 709–722.

Marintchev, A., Edmonds, K.A., Marintcheva, B., Hendrickson, E., Oberer, M., Suzuki, C., Herdy, B., Sonenberg, N., Wagner, G., 2009. Topology and Regulation of the Human eIF4A/4G/4H Helicase Complex in Translation Initiation. Cell 136, 447– 460. doi:http://dx.doi.org/10.1016/j.cell.2009.01.014

McDunn, J.E., Cobb, J.P., 2005. That which does not kill you makes you stronger: a molecular mechanism for preconditioning. Sci. STKE 2005, pe34. doi:10.1126/stke.2912005pe34

Murry, C.E., Jennings, R.B., Reimer, K.A., 1986. Preconditioning with ischemia: a delay of lethal cell injury in ischemic myocardium. Circulation 74, 1124–1136.

Nieuwkoop, P.D., Faber, J., 1956. Normal table of Xenopus laevis (Daudin). A systematical and chronological survey of the development from the fertilized egg till the end of metamorphosis. Norm. table Xenopus laevis (Daudin). A Syst. Chronol. Surv. Dev. from Fertil. egg till end Metamorph. 22.

Nolte, R.T., Wisely, G.B., Westin, S., Cobb, J.E., Lambert, M.H., Kurokawa, R., Rosenfeld, M.G., Willson, T.M., Glass, C.K., Milburn, M. V, 1998. Ligand binding and co-activator assembly of the peroxisome proliferator-activated receptor-γ. Nature 395, 137–143.

Nosjean, O., Boutin, J.A., 2002. Natural ligands of PPARγ:: Are prostaglandin J2 derivatives really playing the part? Cell. Signal. 14, 573–583.

Oberer, M., Marintchev, A., Wagner, G., 2005. Structural basis for the enhancement of eIF4A helicase activity by eIF4G. Genes Dev. 19, 2212–2223.

Panas, M.D., Ivanov, P., Anderson, P., 2016. Mechanistic insights into mammalian stress granule dynamics. J. Cell Biol. 215, 313 LP-323.

Pande, V., Ramos, M.J., 2005. Molecular recognition of 15-deoxy-Δ 12, 14-prostaglandin J 2 by nuclear factor-kappa B and other cellular proteins. Bioorg. Med. Chem. Lett. 15, 4057–4063.

Pereira, M.P., Hurtado, O., Cárdenas, A., Boscá, L., Castillo, J., Dávalos, A., Vivancos, J., Serena, J., Lorenzo, P., Lizasoain, I., 2006. Rosiglitazone and 15-deoxy-Δ12, 14-prostaglandin J2 cause potent neuroprotection after experimental stroke through noncompletely overlapping mechanisms. J. Cereb. Blood Flow Metab. 26, 218–229.

Remm, M., Storm, C.E. V, Sonnhammer, E.L.L., 2001. Automatic clustering of orthologs and in-paralogs from pairwise species comparisons. J. Mol. Biol. 314, 1041–1052.

Rice, P., Longden, I., Bleasby, A., 2000. EMBOSS: the European molecular biology open software suite. Trends Genet. 16, 276–277.

Rogers, G.W., Richter, N.J., Lima, W.F., Merrick, W.C., 2001. Modulation of the helicase activity of eIF4A by eIF4B, eIF4H, and eIF4F. J. Biol. Chem. 276, 30914–30922.

Rozen, F., Edery, I., Meerovitch, K., Dever, T.E., Merrick, W.C., Sonenberg, N., 1990. Bidirectional RNA helicase activity of eucaryotic translation initiation factors 4A and 4F. Mol. Cell. Biol. 10, 1134–1144.

Sali, A., Blundell, T., 1994. Comparative protein modelling by satisfaction of spatial restraints. Protein Struct. by distance Anal. 64, C86.

Scher, J.U., Pillinger, M.H., 2005. 15d-PGJ2: the anti-inflammatory prostaglandin? Clin. Immunol. 114, 100–9. doi:10.1016/j.clim.2004.09.008

Schneider, C.A., Rasband, W.S., Eliceiri, K.W., 2012. NIH Image to ImageJ: 25 years of image analysis. Nat methods 9, 671–675.

Schütz, P., Bumann, M., Oberholzer, A.E., Bieniossek, C., Trachsel, H., Altmann, M., Baumann, U., 2008. Crystal structure of the yeast eIF4A-eIF4G complex: An RNA-helicase controlled by protein–protein interactions. Proc. Natl. Acad. Sci. 105, 9564–9569.

Shibata, T., 2015. 15-Deoxy-Δ12,14-prostaglandin J2 as an electrophilic mediator. Biosci. Biotechnol. Biochem. 79, 1044–1049. doi:10.1080/09168451.2015.1012149

Straus, D.S., Glass, C.K., 2001. Cyclopentenone prostaglandins: new insights on biological activities and cellular targets. Med. Res. Rev. 21, 185–210.

Straus, D.S., Pascual, G., Li, M., Welch, J.S., Ricote, M., Hsiang, C.-H., Sengchanthalangsy, L.L., Ghosh, G., Glass, C.K., 2000. 15-deoxy-Δ12, 14-prostaglandin J2 inhibits multiple steps in the NF-κB signaling pathway. Proc. Natl. Acad. Sci. 97, 4844–4849.

Süel, G.M., Lockless, S.W., Wall, M.A., Ranganathan, R., 2003. Evolutionarily conserved networks of residues mediate allosteric communication in proteins. Nat. Struct. Mol. Biol. 10, 59–69.

Svitkin, Y. V, Gradi, A., Imataka, H., Morino, S., Sonenberg, N., 1999. Eukaryotic initiation factor 4GII (eIF4GII), but not eIF4GI, cleavage correlates with inhibition of host cell protein synthesis after human rhinovirus infection. J. Virol. 73, 3467–3472.

Tan, H.Y., Ng, T.W., 2008. Accurate step wedge calibration for densitometry of electrophoresis gels. Opt. Commun. 281, 3013–3017. doi:http://dx.doi.org/10.1016/j.optcom.2008.01.012

Vieth, M., Hirst, J.D., Brooks, C.L., 1998. Do active site conformations of small ligands correspond to low free-energy solution structures? J. Comput. Aided. Mol. Des. 12, 563–572.

Xu, H.E., Lambert, M.H., Montana, V.G., Parks, D.J., Blanchard, S.G., Brown, P.J., Sternbach, D.D., Lehmann, J.M., Wisely, G.B., Willson, T.M., 1999. Molecular recognition of fatty acids by peroxisome proliferator–activated receptors. Mol. Cell 3, 397–403.

Yang, A.S., Honig, B., 2000. An integrated approach to the analysis and modeling of protein sequences and structures. III. A comparative study of sequence conservation in protein structural families using multiple structural alignments. J. Mol. Biol. 301, 691–711. doi:10.1006/jmbi.2000.3975

Yang, H.-S., Jansen, A.P., Komar, A.A., Zheng, X., Merrick, W.C., Costes, S., Lockett, S.J., Sonenberg, N., Colburn, N.H., 2003. The transformation suppressor Pdcd4 is a novel eukaryotic translation initiation factor 4A binding protein that inhibits translation. Mol. Cell. Biol. 23, 26–37.

Yun, S.J., 2011. Development of a framework for the identification of key factors of biological systems. POSTECH.

